# Epstein-Barr viral product-containing exosome facilitates LIF-associated immunosuppressive polarization of macrophage in nasopharyngeal carcinoma

**DOI:** 10.1101/2024.11.25.625317

**Authors:** Tzu-Tung Liu, Yun-Hua Sui, Shih-Sheng Jiang, Fang-Yu Tsai, Kai-Ping Chang, Ngan-Ming Tsang, Meng-Hsin Lee, Chen-Han Huang, Shu-Chen Liu

**Affiliations:** Department of Biomedical Sciences and Engineering, National Central University, Taiwan; Institute of Cancer Research, National Health Research Institutes, Taiwan; Department of Otolaryngology-Head & Neck Surgery, Linkou Chang Gung Memorial Hospital, Taiwan; Department of Radiation Oncology, China Medical University Hsinchu Hospital, Zhubei City, Hsinchu County, Taiwan

**Keywords:** Nasopharyngeal carcinoma, Epstein-Barr virus, exosome, tumor-associated macrophages, leukemia inhibitory factor, immune evasion, tumor microenvironment

## Abstract

Epstein-Barr virus (EBV)-associated nasopharyngeal carcinoma (NPC) is characterized by a highly infiltrated, yet immunosuppressive, tumor microenvironment (TME) in which tumor-associated macrophages (TAMs) play a central role in tumor progression. In this study, we identified exosome-mediated delivery of the EBV oncoprotein LMP1, which acts via induction of leukemia inhibitory factor (LIF), as a key mechanism driving immunosuppressive macrophage polarization. Immunohistochemistry revealed that, in addition to tumor cells, LIF is primarily expressed by stromal TAMs and correlates with poor prognosis. We demonstrated that LMP1-containing exosomes are internalized by macrophages, leading to NF-κB-dependent upregulation of LIF and polarization toward an immunosuppressive M2d-like phenotype. These macrophages exhibit impaired anti-tumor activity and promote tumor proliferation and vascular dissemination in a LIF-dependent manner. Single-cell transcriptomic analyses of exosome-treated NPC biopsies revealed transcriptional reprogramming across tumor, myeloid, and lymphoid compartments impacting epithelial-mesenchymal transition (EMT); extracellular matrix (ECM) remodeling in tumor cells; immunosuppressive signaling (e.g., CD163 anti-inflammatory and VEGF signaling in macrophages); and enrichment of immune checkpoints and pathways, including PD-1, TGFβ-SMAD, and NF-κB-associated pathways in T cells. Exosome treatment increased expansion of CCL18^+^-TAMs and expression of CTLA4 in regulatory T cells and exhausted T cells. Immunohistochemistry further confirmed a positive correlation between expression of LIF and CTLA4 in NPC tumors and an inverse correlation between LIF and GZMB/CD8A. These findings define an exosome-driven LMP1-LIF axis that orchestrates immune suppression in NPC and suggest potential targets for restoring anti-tumor immunity.

## Introduction

Non-keratinizing nasopharyngeal carcinomas (NPCs) are strongly associated with Epstein-Barr virus (EBV) infection, which plays a crucial role in remodeling the immune microenvironment of NPC (1). Despite heavy infiltration of NPC lesions by immune cells, including T and B lymphocytes, macrophages, neutrophils and dendritic cells, the anti-tumor activity within these lesions is often ineffective.

EBV and the host immune system engage in a multifaceted interplay. EBV modulates interactions between tumors and stromal cells, promoting the recruitment and activation of immunosuppressive populations such as regulatory T cells (Tregs), M2-like macrophages, and myeloid-derived suppressor cells (MDSCs) (2,3). Several EBV-encoded products, including LMP1, LMP2, BZLF1, EBNA1 and EBERs (3–5), contribute to immune evasion by inducing immunoregulatory factors such as IL-10, TGFB1 (transforming growth factor beta 1), IL-1Ra, GM-CSF, IL-8, and VEGF (vascular endothelial growth factor) (5–7). These factors reshape the TME, promoting immune suppression and tumor progression. In addition to direct signaling, EBV-positive NPC cells influence the TME through the release of exosomes. These vesicles contain viral components such as LMP1, FAPα and miR-BART15-3p (8–10), which can alter immune cell behavior, support tumor growth, and facilitate viral persistence. TAMs are enriched in NPC tumors. In EBV-infected NPCs, TAMs frequently present an immunosuppressive type, but the upstream signals driving this polarization in NPC are not fully defined. Although exosomal communication is increasingly recognized as a mechanism of intercellular signaling within the TME, the precise contribution of EBV product-containing exosomes to macrophage polarization and immune suppression in NPC remains unclear.

Leukemia inhibitory factor (LIF) is a member of the IL-6-type cytokine family. Signaling downstream of LIF is mediated by interactions with the LIF receptor (LIFR) and Gp130, triggering signaling cascades such as the JAK/STAT3, PI3K, mTOR/p70S6K1, and ERK1/2 signaling pathways to regulate cell proliferation, survival, differentiation, and other biological processes (11). LIF can be secreted by many cell types, including macrophages, fibroblasts and cancer cells, and its production can be induced by the inflammatory factors NF-κB, TNF-α and IL-1, among others (12). LIF promotes tumor progression, metastasis and chemoresistance in many solid tumors (13–19), but also has a pivotal role in specific regulation of adaptive immunity (20). High levels of LIF can be released by activated human Tregs (20), whereas it has been shown that LIF activity is shut down in Th17 cells (21). Moreover, there is evidence that Gp130 and LIFR expression are very low in un-stimulated, naive T cells, but these cells undergo a conversion to a Gp130^high^LFIR^high^ phenotype within 48□hours of activation (22). In addition, LIF favors acquisition of an M2 phenotype by macrophages and recruitment of myeloid-derived suppressor cells into the TME (23–25). It has also been demonstrated that tumors expressing high levels of LIF tend to be infiltrated with TAMs (26). In NPC, however, the cellular origin and regulatory mechanisms of LIF within the TME have not been systematically investigated. Notably, EBV-encoded LMP1 has been shown to induce LIF production in NPC cells (27), but whether LMP1 influences macrophage LIF expression via exosomal delivery is unknown.

Given the prominent presence of EBV-infected cells and macrophages in NPC, and the established role of LIF in modulating immune responses, we hypothesized that exosome-mediated delivery of LMP1 drives macrophage polarization through induction of LIF. To test this, we investigated the spatial distribution of LIF in the NPC TME, the transfer of LMP1 to macrophages via tumor-derived exosomes, and consequent effects on macrophage phenotype and tumor-immune interactions. By integrating immunohistochemistry (IHC), single-cell transcriptomics and functional analyses, our study identifies the exosomal LMP1-LIF axis as a key driver of macrophage-mediated immune suppression in NPC. We show that EBV-LMP1-containing exosomes reprogram macrophages toward a pro-tumoral type, which promotes cancer cell proliferation, dissemination, and T cell dysfunction. These effects are mediated through NF-κB-LIF signaling. Targeting this exosome-mediated crosstalk may represent a promising strategy for reversing immune suppression and improving outcomes in NPC.

## Materials and Methods

### Clinical samples

A total of 239 pretreatment formalin-fixed paraffin-embedded (FFPE) NPC tissue samples were collected from the tissue bank of Chang Gung Memorial Hospital, Taiwan. All tumors were histologically confirmed by pathologists. All patients completed treatment with either radiotherapy or concurrent chemoradiotherapy at Chang Gung Memorial Hospital. All individuals were followed for more than 5 years post treatment. Ethical approval was granted by the Institutional Review Board of Chang Gung Memorial Hospital, Taiwan (IRB104-9216B, IRB202202095B0). Informed consent was waived due to the retrospective nature of the study.

### Cell culture

HK1 EBV (EBV^+^-human NPC cell line) (28), B95-8 (EBV^+^-lymphoblastoid cell line) (29), and BM1 (human NPC cell line) (30) cells were cultured in RPMI-1640 medium supplemented with 10% fetal bovine serum (FBS) (Peak) and 1% penicillin-streptomycin (P/S) (Corning). NPC-TW06 cells (31) cells were cultured in DMEM medium supplemented with 10% FBS and 1% P/S. Primary PBMC monocytes (Lonza Bioscience) were cultured in α-MEM medium supplemented with 10% FBS and 1% P/S. Primary cancer cells were cultured in defined serum-free keratinocyte medium (PromoCell). The authenticity of HK1EBV, BM1, NPC-TW06, and B95-8 cells was confirmed by 16-core short tandem repeat (STR) profiling (Bioresource Collection and Research Center, Taiwan).

### Isolation and labeling of exosomes

Exosomes were isolated as previously described (9). Briefly, culture supernatants from HK1 EBV cells grown for 2 days in RPMI-1640 with 1% EV-depleted FBS were collected. Exosomes were isolated using ExoQuick-TC (System Biosciences) and resuspended in cold PBS, aliquoted and stored at −80°C until use. For tracking, exosomes were labeled with ExoGlow™-Protein EV Labeling Kit (System Biosciences). For western blotting, exosome pellets were lysed in 1% NP-40 lysis buffer and stored at − 80 °C until use.

### Identification of EBV-encoded products in exosomes

EBV-derived products in exosomes were identified using LC/ESI-MS/MS (Academia Sinica, Taiwan). Briefly, 10□mg exosome lysates were reduced with dithiothreitol (DTT), alkylated with 55□mM iodoacetamide in 50□mM ammonium bicarbonate, digested with trypsin, and desalted using ZipTip □18. 0.5□mg of protein digests were analyzed using a 90-minute LC gradient on Orbitrap Elite MS (Thermo Fisher Scientific). Raw data were processed with Proteome Discoverer and searched against the Uniprot EBV database using MSFragger (32). Peptides with > 99% probability were accepted.

### Transmission electron microscopy (TEM) analysis of exosomes

TEM was performed as previously described (9). Fresh exosomes were captured using anti-CD9 antibody-conjugated Dynabeads (Invitrogen). Beads were blocked with 1% BSA/PBS, washed, and applied to mesh copper grids. After an 8-minute incubation, grids were washed with ddH□O. Exosomes were visualized using a JEOL JEM2000FX transmission electron microscope operating at 200□kV.

### Differentiation of human PBMC-derived macrophages

PBMC monocytes were cultured in α-MEM supplemented with GM-CSF for 1.5□hours, then washed and incubated in α-MEM supplemented with 10% FBS. On day 7, macrophages were treated as follows: 50□ng/ml IFNg and 10□ng/ml LPS (M1-like), 20□ng/ml IL-4 (M2-like), 5□ng/ml TGFβ (M2c), 10□ng/ml IL-6 (M2d), or 30□μg/ml exosomes (exosome-stimulated). Untreated macrophages were defined as M0 (Mock). Medium was replaced, and stimulation was repeated on day 9. Macrophages were considered fully differentiated on day 11 or day 12.

### Zebrafish embryonic xenograft tumor model

We used Tg (fli1a:EGFP) y1 transgenic zebrafish (Core Facility, NHRI, Taiwan) to assess vascular dissemination (33). Approximately 300 NPC-TW06_LifeAct-RFP cells were injected into the yolk sac within one hour post fertilization. Tumor growth and dissemination were monitored from day 5 to day 8 using fluorescence microscopy (Olympus IX83). Quantification was performed using CellSens software. Experiments followed the guidelines of the Institutional Animal Care and Use Committee (IACUC) of National Health Research Institutes (NHRI), Taiwan.

### Macrophage phagocytic assay of NPC spheroids

NPC spheroids were generated using AggreWell™800 24-well plate (STEMCELL Technologies). NPC-TW06_LifeAct-RFP cells were seeded and grown for 24 hours to allow spheroid formation. Macrophages pretreated with vehicle or 30 μg/ml exosomes derived from HK1EBV, and/or 1 μg/ml sLIFR, were added at a 1:1 effector-to-target ratio. Macrophages were pre-labeled with CellTracker, Thermo). After 48 hours of coculture, spheroids were imaged using Olympus IX83 microscopy and analyzed with CellSens software.

### Transfection

Macrophages were transfected with full-length LMP1 plasmids (pCMV-FL-LMP1) (34,35) or vector control using Fugene HD (Promega). Cell lysates were harvested at 24 and 36 hours. HK1EBV cells were transfected with EBV-LMP1 or control siRNA (MDBio, Taiwan) using RNAiMAX reagent (Invitrogen). siRNA sequences are: LMP1 siRNA-1: 5′-GACCUCAUCCUGCUCAUUATT-3′. LMP1 siRNA-2: 5′-GGACCCUGACAACACUGAUTT-3′. Negative control: 5′-UUCUCCGAACGUGUCACGUTT-3′.

### Generation of 3**D**-single cell expression libraries

Fresh pretreatment NPC biopsies were obtained from Chang Gung Memorial Hospital. Samples were immediately washed and trimmed into small pieces, followed by enzymatic treatment and mechanical dissociation (Miltenyi Biotech). Cell suspensions were stringently washed and filtered through a 40 μm cell strainer. Cell viability was assessed by Calcein-AM/PI double staining. Cells (2×10^6^ cells/well in 6-well plate) with viability greater than 90% were treated with exosomes or vehicle for 2 hours at 37°C, then loaded into microfluidic chips to generate gel beads-in-emulsion (GEMs). Reverse transcription and cDNA amplification were performed using Chromium™ single cell 3’ v3.1 reagent kit (10x Genomics). Libraries were sequenced on a NovaSeq 6000 Sequencing System (Illumina).

### Processing of single-cell RNA sequencing (ScRNA-seq) data

Raw BCL files were processed using Cell Ranger (v6.1.2). Data were demultiplexed, converted to FASTQ, and aligned to the human genome (GRCh38). Seurat R package (version 5.1.0) (36) was used for quality control, normalization, dimension reduction, clustering, and differential expression (DE) analysis. Cells with fewer than 200 or more than 8000 feature counts or over 10% mitochondrial genes were excluded. Doublets were further removed using DoubletFinder (version 2.0.3) (37).

### Functional analysis of scRNA-seq data

UCell R package (version 2.6.2) (38) was used to calculate signature scores (Supplementary Table S1) and GSVA R package (version 1.50.5) (39) was used to calculate enrichment scores based on gene sets from the Molecular Signatures Database (MSigDB). Cell-cell communication analysis was performed using NicheNet (version 2.1.5) (40). Cancer cells and TAMs were designed as sender cells, and T cells as receiver cells. Genes expressed in more than 5% of cells were considered “expressed genes”. Exosome-upregulated DEGs associated with co-inhibitory T cell function were designated as “genes of interest”. Ligands were prioritized based on the area under the precision-recall curve (AUPR) using NicheNet’s predict_ligand_activities.

### Study approval

Ethical approval was granted by the Institutional Review Board of Chang Gung Memorial Hospital, Taiwan (IRB104-9216B, IRB202202095B0).

### Antibodies

Primary antibodies used in western blotting include: CD9 (Abcam, ab92726), HSP70 (System Biosciences, EXOAB-HSP70A), Calnexin (Cell Signaling, #2679), β-actin (Sigma, A5060), CD81 (Santa Cruz Biotechnology, sc-166029), CD206 (Abcam, ab64693), DC-SIGN (BD Biosciences, MAB161), VEGFA (GeneTex, GTX637405), CD86 (Cell Signaling, #91882), CD80 (ABclonal, A19025), GAPDH (Abcam, ab9484), LIF (Abcepta, AP6981C), phospho-IκBa (Abcam, ab97783), and IκBa (Santa Cruz Biotechnology, sc-371), LMP1 (S12; purified from hybridoma culture supernatants). Primary antibodies for IHC or ICC include: EBV-encoded LMP1 (S12; purified from hybridoma culture supernatants), LIF (Abcam, ab135629), EBV early antigen D (EaD; Abcam, ab30541), and CD68 (Abcam, ab213363).

### Statistical analysis

Statistical analysis was conducted using R and GraphPad Prism 10. Kaplan-Meier survival and log-rank tests were used for survival comparison. Chi-square tests were used to evaluate difference in IHC scores. Spearman’s correlation test was used for IHC-based correlation analysis The Mann-Whitney U test and Student’s t-test were used to assess exosome-mediated effects. All statistical tests were two-sided and P values < 0.05 considered statistically significant.

### Data availability

All data supporting the findings of this study are available within this published article, its supplemental data files, and from the corresponding author upon reasonable request. The raw and processed single-cell data have been deposited in the Gene Expression Omnibus (GEO) under accession number GSE280127. The bulked RNA sequencing data of macrophages have been deposited in GEO (GSE282685) as well.

## Results

### Elevated LIF levels in the NPC tumor microenvironment correlate with poor prognosis and macrophage localization

LIF is increasingly recognized for its pivotal role in tumorigenesis and immune modulation (23,41,42). Our IHC analysis predominantly detected LIF immunoreactivity in NPC tumor cells and immune cells within the TME (Fig. 1A). Notably, LIF-positive immune cells were mainly located in stromal areas or at tumor borders, although some were present within tumor nests. Quantification of LIF expression within the NPC TME, encompassing both tumor and immune cells, revealed significantly higher levels in patients with distal metastasis (P = 0.0019; Fig. 1B). Survival analyses showed that patients with elevated TME LIF levels had poorer recurrence-free survival (P = 0.04; Fig. 1C) and metastasis-free survival (P = 0.016; Fig. 1D) compared with those with lower levels. To further investigate the cellular source of LIF, we examined CD68 and LIF expression in consecutive NPC biopsy sections. We observed partial colocalization of LIF-positive immune cells with CD68-positive macrophages in stromal areas (Fig. 1E, NPC#1), whereas such colocalization was absent in macrophages infiltrating tumor nests (Fig. 1E, NPC#2). Quantitative analyses revealed that immune cells in the stroma expressed significantly higher levels of both LIF (79%) and CD68 (75%) compared with those in tumor nests (P < 0.0001; Fig. 1F). Only approximately 7.5% of NPC cases exhibited detectable (score ≥ 15) LIF-positive immune cells in tumor nests, generally with weak immunoreactivity. In contrast, CD68-positive macrophages, typically lacking LIF expression, were found to infiltrate about 38% of tumor nests. Correlation analyses revealed a strong association between LIF and CD68 expression in the stroma (r = 0.5012, P = 0.0003; Fig. 1G), but no significant correlation within tumor nests (r = 0.0999, P = 0.4855; Fig. 1H). Double-immunofluorescence staining confirmed that tumor-infiltrating macrophages were larger and lacked LIF expression (Fig. 1I, upper panels, NPC#1), whereas LIF-positive macrophages were localized to stromal regions (Fig. 1I, lower panels, NPC#2). A quantitative analysis of 5545 CD68-positive macrophages from six NPC cases showed that ∼22% of stromal macrophages were LIF-positive, whereas only ∼3% of tumor-infiltrating macrophages expressed LIF (Fig. 1J). Together, these data indicate that LIF expression is associated with stromal macrophages rather than tumor nest-infiltrating macrophages, suggesting that LIF may serve as a marker of immunosuppressive TAMs in the NPC TME.

**Figure 1.**
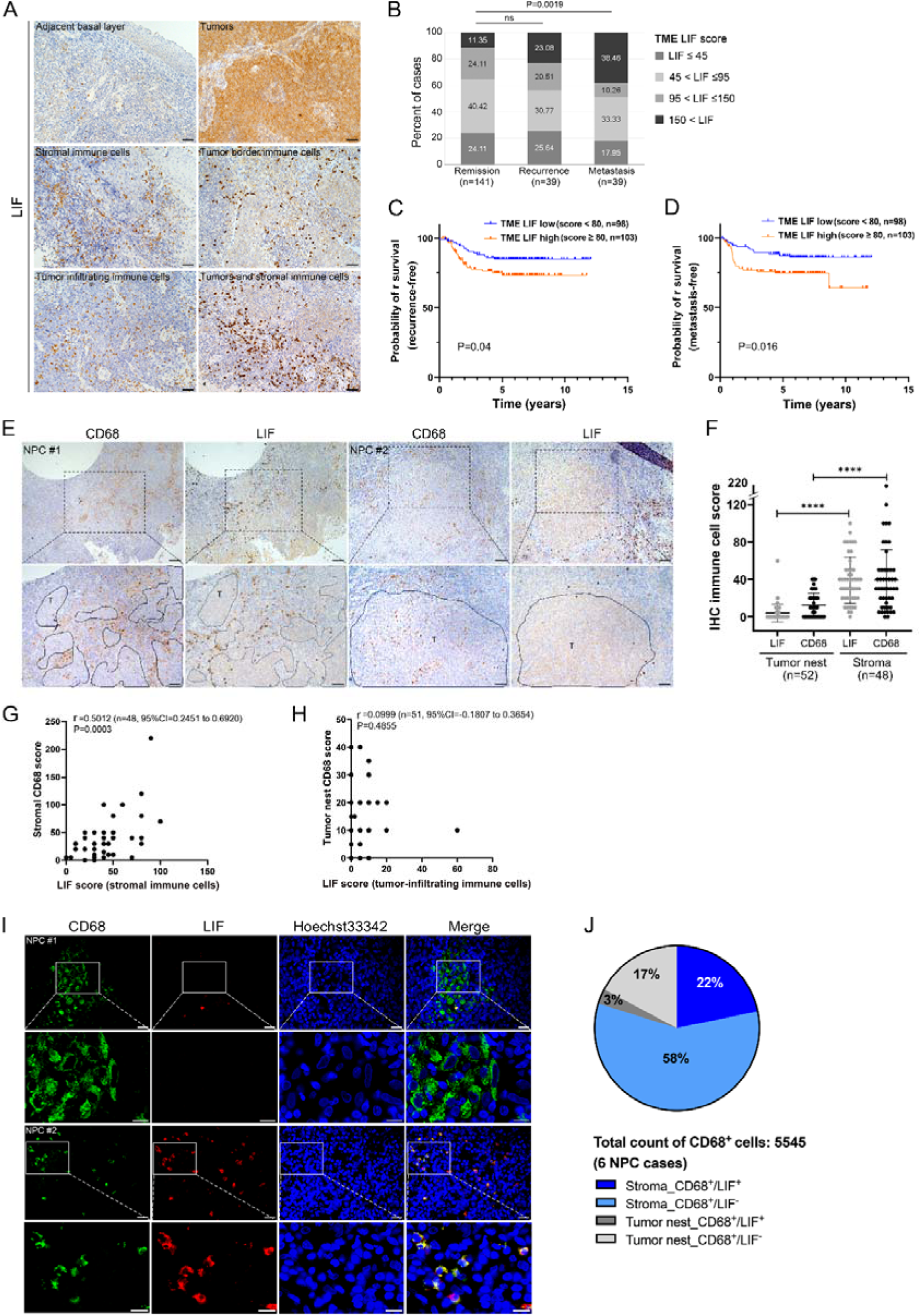
Elevated LIF levels in the NPC TME are associated with poor prognosis and macrophage localization. (A) Representative IHC images of LIF expression in NPC tumor cells and immune cells within the TME. Scale bars, 50 μm. (B) Quantification of overall LIF expression in the TME of NPC patients (Fisher’s exact test). (C and D) Kaplan-Meier survival analysis of NPC patients based on TME LIF levels showing recurrence-free survival (C) and metastasis-free survival (D). (E) Representative IHC images of CD68 and LIF in consecutive NPC sections. Black areas (bottom panels) indicate tumor nests. Scale bars, 100 μm and 50 μm (enlarged). (F) Quantification of LIF- or CD68-positive immune cells in stromal areas versus tumor nests (Mann-Whitney U test). (G and H) Correlation between immune cell LIF and CD68 expression scores in stromal areas (G) and tumor nests (H) (Spearman’s correlation test). (I) Representative double-immunofluorescence images of CD68 (green) and LIF (red) expression in macrophages within tumor nests (NPC#1) and stromal regions (NPC#2). Scale bars, 20 μm and 10 μm (enlarged). (J) Quantification of LIF-positive CD68^+^ macrophages in stromal and intratumoral regions across six NPC cases (total of 5545 macrophages).

### Detection of EBV-encoded LMP1 in macrophages within NPC tissue sections and NPC-derived exosomes

Given that EBV-encoded LMP1 upregulates LIF production in NPC (27), we speculated that LIF expression in TAMs might reflect exosomal transfer of LMP1 to TAMs in the NPC microenvironment. We examined the expression of EBV-encoded genes, including EBV-encoded small RNAs (EBERs), early antigen D (EaD) and LMP1, in consecutive NPC sections and found that, in addition to its presence in tumor cells, LMP1 immunoreactivity was more frequently detected in stromal cells than was EBERs or EaD immunoreactivity (Fig. 2A). Expression of LMP1 was not only detected in the normal basal layer and tumor nests, but also in surrounding immune cells (Fig. 2B). Immunofluorescence staining further confirmed the presence of LMP1 in CD68□ macrophages in NPC biopsies (Fig. 2C). To assess whether the LMP1 positivity in TAMs is attributable to exosomal delivery, we isolated exosomes from EBV-positive NPC cells (HK1EBV) (28) and the EBV-positive lymphoid cell lines, B95-8 (29) and AKATA (43). Transmission electron microscopy (TEM) characterization of exosomes captured by magnetic beads conjugated with anti-CD9 antibodies revealed a vesicle diameter of ∼100 nm (Fig. 2D). HK1EBV-derived exosomes were taken up by peripheral blood mononuclear cell (PBMC)-derived macrophages, as determined using exosomes prelabeled with ExoGlow (green) (Fig. 2E). Stimulation of macrophages with these exosomes led to increased expression of EBV genes, including *EBER1, LMP1, BZLF1* and *BXLF1* (which encodes EBV thymidine kinase, TK) as measured by quantitative reverse transcription-polymerase chain reaction (qRT-PCR) (Fig. 2F). Western blot analyses further confirmed the presence of LMP1 in exosomes from both HK1EBV and B95-8 cells, but not those from EBV-negative NPC cells (BM1) (Fig. 2G). Expression of other viral genes, such as *BPLF1* and *BXLF*, was also detected in exosomes derived from EBV^+^ cancer cells, as characterized by qualitative mass spectrometry (MS) (Supplementary Table S3). Moreover, LMP1 protein was detected in CD68□ macrophages treated with exosomes from EBV□ cell lines (Fig. 2H). These results suggest that LMP1 can be delivered to and expressed in macrophages via tumor-derived exosomes.

**Figure 2.**
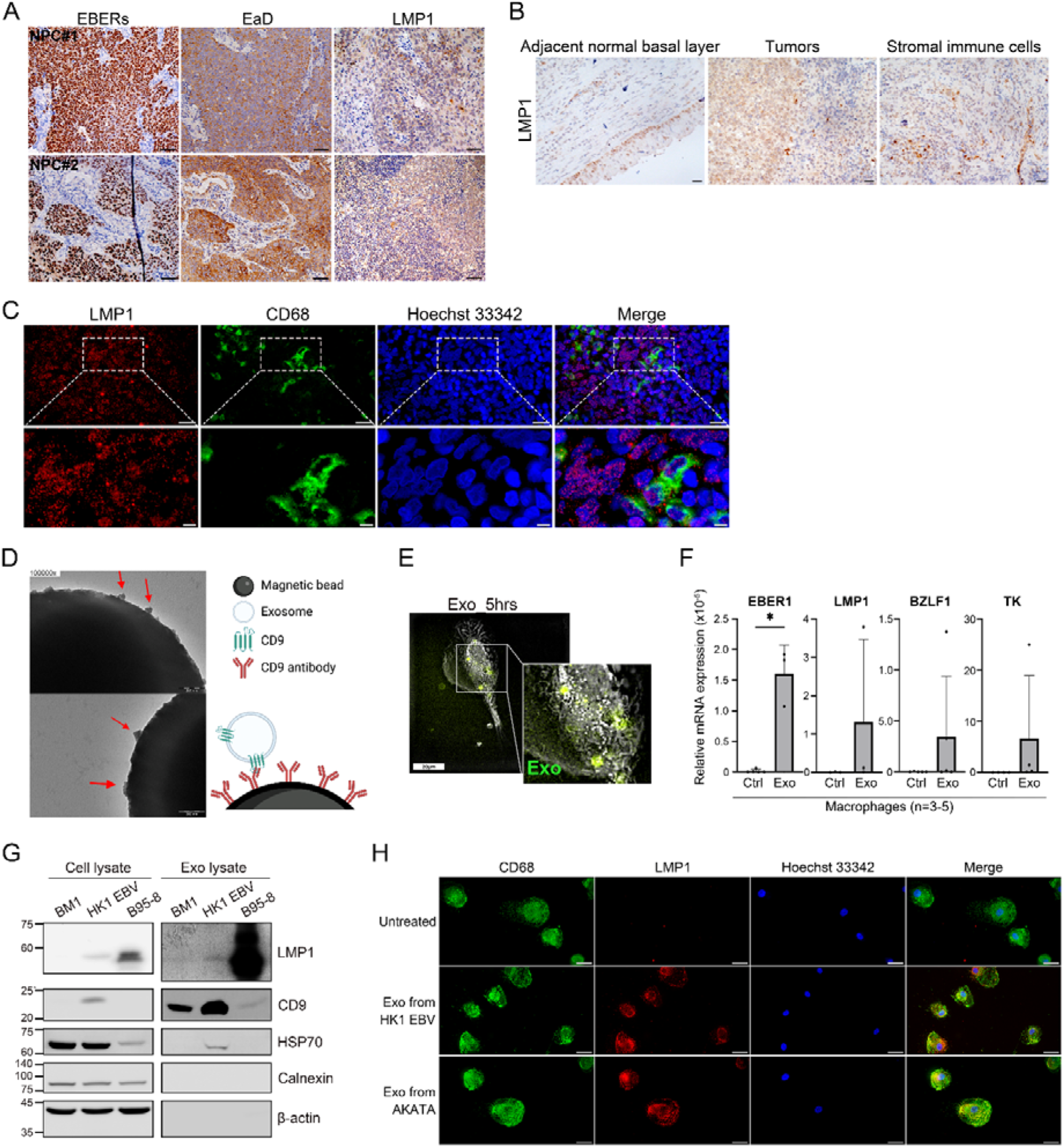
Presence of EBV-encoded LMP1 in macrophages within NPC tissue sections and NPC-derived exosomes. (A) Detection of EBERs, EaD, and LMP1 in consecutive NPC tissue sections. Scale bars, 100 μm. (B) Representative images of IHC staining for LMP1 in the normal basal layer, tumor nests, and immune cells. Scale bars, 50 μm. (C) Representative images of double-immunofluorescence staining for LMP1 (red) and CD68 (green) in NPC tissue sections. Nuclei were counterstained with Hoechst 33342. Scale bars, 20 μm and 10 μm (enlarged). (D) TEM images of exosomes captured using CD9-coated immunomagnetic beads. Arrows indicate captured exosomes. Scale bars, 200 nm. (E) Uptake of ExoGlow-labeled exosomes (green) by macrophages. Images were captured 5 hours post treatment. Scale bars, 10 μm. (F) Relative expression of EBV genes (*EBER1*, *LMP1*, *BZLF1*, and *BXLF1*(which encodes EBV thymidine kinase, TK in PBMC-derived macrophages stimulated for 3 hours with exosomes from HK1EBV cells. Expression levels were normalized to β-actin (ACTB) (*P < 0.05). (G) Western blot analysis of LMP1 and exosomal proteins in cell lysates and exosomes from EBV-negative NPC cells (BM1), EBV-positive NPC cells (HK1EBV), and B95.8 cells. Samples were collected after 2 days of culture. (H) Representative images of double-immunofluorescence staining for CD68 (green) and LMP1 (red) in PBMC-derived macrophages treated with exosomes from HK1EBV or EBV□ AKATA cells for 3 hours. Nuclei were counterstained with Hoechst 33342. Scale bars, 20 μm.

### EBV product-containing exosomes induce immunosuppressive macrophages, which promote NPC cell proliferation

To investigate whether EBV-derived exosomes influence macrophage polarization, we differentiated PBMC-derived monocytes in the presence of HK1EBV exosomes. The resulting macrophages displayed a distinct morphology compared to classical M1 or M2 subtypes. By day 9 post differentiation, M1-like macrophages exhibited a rounded and flattened phenotype, whereas M2-like and M2d macrophages, the latter of which are a predominantly pro-cancer subtype that makes a major contribution to the TAM population, were more elongated. The morphology of exosome-treated cells resembled that of M2d-like macrophages (Fig. 3A). Transcriptomic profiling by bulk RNA sequencing and principal component analysis (PCA) revealed that exosome-treated macrophages clustered separately from classical polarization subtypes, but were closely aligned with M2d-like macrophages (Fig. 3B). A heatmap analysis showed comparable interferon signaling, pro-inflammatory response, and immune suppression expression profiles between exosome-treated and M2d-like macrophages (Fig. 3C). Notably, exosome treatment upregulated immunosuppressive markers, including IL-10, PTGER4, PD-L1 (CD274), PDCD1LG2 (PD-L2), and TGFB1. qRT-PCR analysis revealed that the expression patterns of M2d-associated genes, including *CD206*, *IL10*, *ARG1* and *HLA-DRA*, were similar between exosome-treated and M2d-like macrophages (Fig. 3D). Western blot analysis showed increased expression of CD206 and VEGF, and reduced expression of CD86 and CD80, consistent with an immunosuppressive phenotype (Fig. 3E). Functionally, exosome-treated macrophages exhibited impaired phagocytic activity against primary cancer cells, which appeared to escape macrophage-mediated attack (Supplementary Video-1). This contrasts with phagocytic events by M0 macrophages (control group), which continuously attacked cancer cells and eventually engulfed them ∼40 hours after their addition (Supplementary Video-2). These findings suggest that EBV product-containing exosomes facilitate cancer immune evasion. To examine the effect of these macrophages on cancer growth, we treated NPC cells with macrophage-conditioned medium (CM). Results of real-time proliferation assays (Fig. 3G) and EdU incorporation analysis (Fig. 3H and I) showed that CM from exosome-treated macrophages (Exo-Tx Mφ^CM^) significantly enhanced NPC cell proliferation compared with cells treated with M0 Mφ^CM^ or mock controls (P < 0.0001). Collectively, these results indicate that EBV product-containing exosomes promote polarization of immunosuppressive macrophages, which, in turn, support NPC cell proliferation.

**Figure 3.**
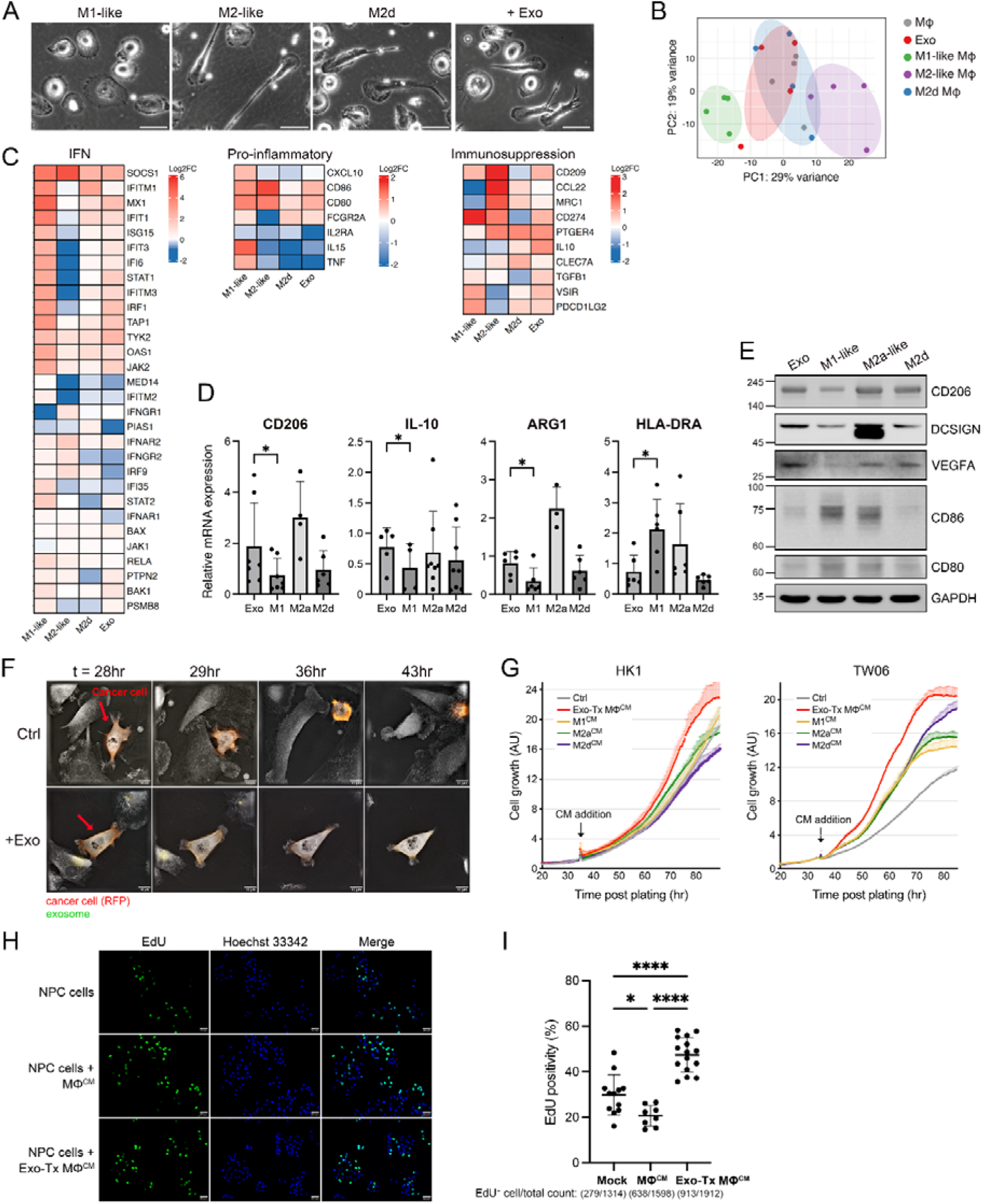
EBV product-containing exosomes induce immunosuppressive macrophage polarization and promote NPC cell proliferation. (A) Representative morphology of macrophages differentiated from PBMCs using established polarizing factors or HK1EBV-derived exosomes. Images were captured on day 5 post treatment. Scale bars, 50 μm. (B) Principal component analysis (PCA) of bulk RNA-seq data from macrophages harvested on day 12 post differentiation. (C) Expression heatmaps of genes involved in interferon (IFN), pro-inflammatory, and immunosuppressive signaling in subtypes of macrophages. Data are presented as log_2_ fold changes relative to M0 macrophages. Red indicates upregulation and blue indicates downregulation. (D) Quantitative RT-PCR analysis of *CD206*, *IL-10*, *ARG1*, and *HLA-DRA* expression in macrophage subtypes. (E) Western blot analysis of polarization markers in PBMC-derived macrophages. GAPDH was used as a loading control. (F) Phagocytic analysis of PBMC-derived macrophages cocultured with primary cancer cells (red arrows). Time zero (t=0) indicates the addition of cancer cells. Scale bars, 50 μm. (G) Real-time proliferation analysis of NPC cells treated with CM from macrophage cultures. Arrows indicate the time of CM addition. Data are presented as means and SD of triplicate experiments. (H) EdU incorporation assays in NPC-TW06 cells treated with macrophage (Mφ) CM or control medium for 18 hours. EdU (green) was added for the final 4 hours. Nuclei were counterstained with Hoechst 33342. Scale bars, 50 μm. (I) Quantification of EdU incorporation shown in (H). *P < 0.05, ****P < 0.0001 (Bonferroni’s multiple comparison test).

### LMP1-containing exosomes upregulate LIF, which mediates pro-tumor macrophage polarization

EBV-encoded LMP1, the major EBV oncoprotein in NPC, has been reported to induce LIF production in NPC cells (27), thereby promoting NPC progression and contributing to poorer prognosis. We investigated whether uptake of LMP1-containing exosomes by macrophages induces LIF expression and contributes to macrophage polarization. We found that exosome exposure led to increased expression of LIF and phospho-IκB (S32), together with upregulation of macrophage polarization markers (Fig. 4A and B). These effects were observed upon addition of exosomes during differentiation (Fig. 4A) as well as with short-term stimulation of differentiated macrophages with exosomes (Fig. 4B). To determine whether these effects depend on exosomal LMP1, we depleted LMP1 in HK1EBV cells prior to exosome isolation. As shown in Figure 4C, exosomal LMP1 protein was detected in control exosomes, but was clearly decreased following LMP1 knockdown. Consistent with this, exosomes from LMP1-depleted cells did not enhance expression of LIF or phospho-IκB in recipient macrophages. Furthermore, ectopic expression of LMP1 in macrophages increased LIF and phospho-IκB expression (Fig. 4E), confirming the role of exosomal LMP1 (Fig. 4D) in regulating LIF production and macrophage polarization. To assess the functional consequences of LIF on NPC cells, we blocked LIF-mediated effects by co-treating cells with CM and soluble LIF receptor (sLIFR), a LIF antagonist. Results of these experiments demonstrated that Exo-Tx Mφ^CM^-enhanced NPC cell proliferation was suppressed by sLIFR in both HK1EBV and HK1 cells (Fig. 4F). To assess the impacts of Exo-Tx Mφ^CM^ on cancer invasive behavior, we used a zebrafish xenograft model, injecting NPC-TW06_LifeAct-RFP cells into the yolk sac of embryos of Tg (fli1a:EGFP) y1, an EGFP-positive endothelial transgenic zebrafish line, and monitoring the growth and dissemination of tumor-like structures for 7 days (Fig. 4G). As shown in Figure 4H, tumor-like structures were observed at the yolk sac, where the cancer cells were injected, and at distal sites near blood vessels. Statistical analyses demonstrated that, compared with the control group, zebrafish in the Exo-Tx Mφ^CM^ group exhibited enhanced NPC vascular dissemination, an effect that was significantly inhibited in the presence of sLIFR (Fig. 4I). We further evaluated the anti-cancer immunity of macrophages in a 3D cancer spheroid model in which fluorescently labeled macrophages (M□_GFP) were added to 1-day NPC-TW06_LifeAct-RFP spheroids (NPC_RFP) and cocultured for an additional 2 days. These experiments showed that the growth of NPC spheroids was enhanced by coculture with exosome-treated macrophages, an effect that was suppressed by sLIFR-mediated targeting of LIF (Fig. 4J and K). These results indicate that exosomal LMP1 promotes pro-tumoral macrophage polarization by inducing LIF expression, suggesting that the exosomal LMP1-LIF axis mediates tumor-immune crosstalk in the NPC TME.

**Figure 4.**
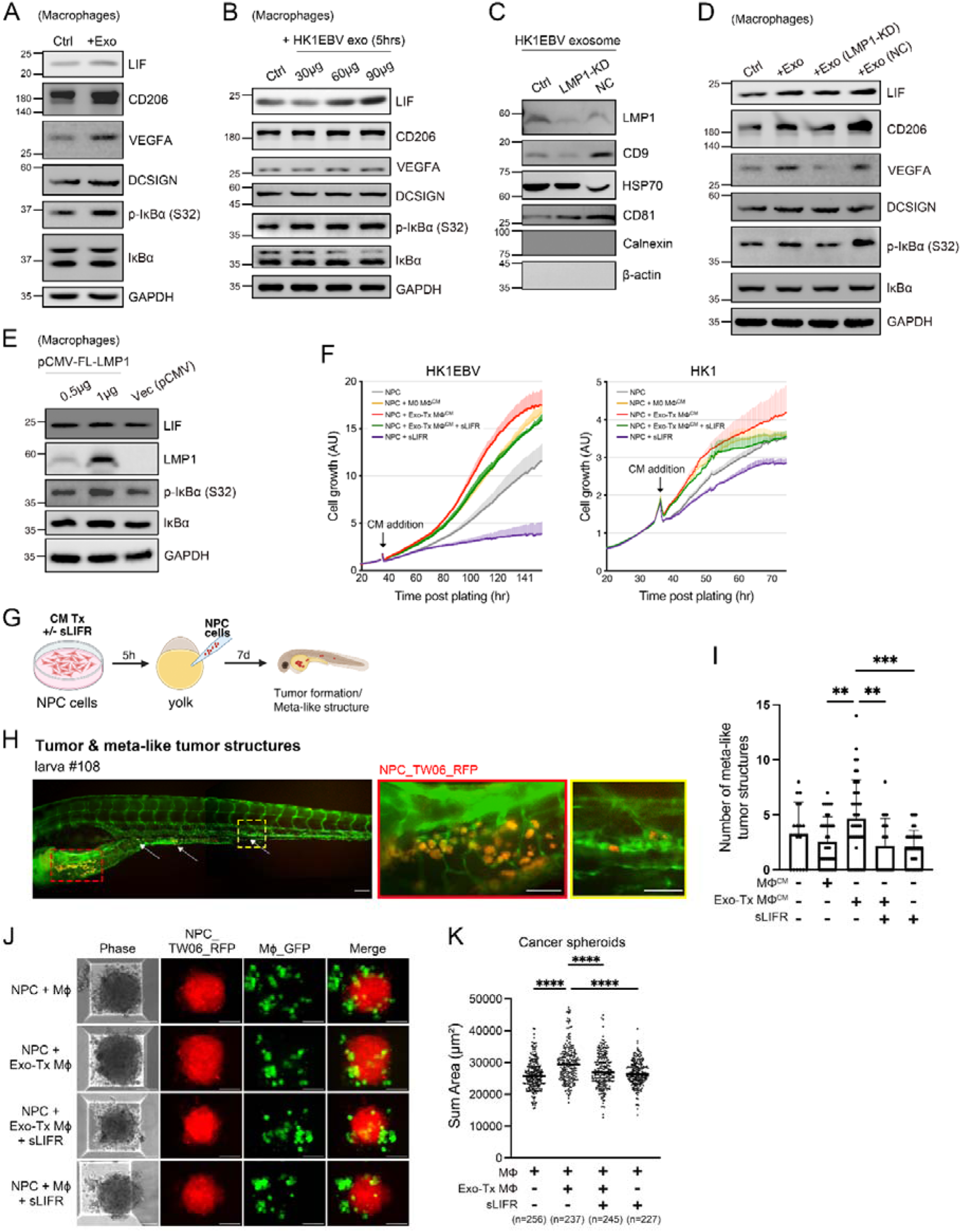
Exosomal LMP1 induces pro-tumoral macrophage polarization through LIF production. (A and B) Western blot analysis of macrophage differentiation markers, phospho-IκB (S32), and LIF in PBMC-derived macrophages treated with exosomes or control medium. HK1EBV-derived exosomes (50 μg/ml) were added on day 3 during differentiation (A) or applied to differentiated macrophages for 5 hours with increasing concentrations of exosomes (B). GAPDH was used as a loading control. (C) Western blot detection of LMP1 and exosome markers in exosomes isolated from HK1EBV cells transfected with LMP1 siRNA, NC siRNA, or mock control. Exosomes were isolated 2 days post transfection. (D) Western blot analysis of macrophages treated with exosomes from HK1EBV cells transfected with either LMP1 siRNA (LMP1-KD) or control siRNA (NC). Protein lysates were harvested 2 days post transfection. (E) Western blot detection of LIF and phospho-IκB (S32) expression in macrophages transfected with either full-length LMP1 or a control vector. Cell lysates were harvested 36 hours post transfection. (F) Real-time growth analysis of NPC cells treated with CM from macrophage cultures in the presence of sLIFR or vehicle (PBS). The black arrow indicates the time of CM/sLIFR addition. Data are presented as mean and SD of triplicate experiments. (G) Schematic depiction of the zebrafish embryonic xenograft model. NPC-TW06 cells expressing LifeAct-RFP were injected into the yolk sac of embryos of Tg (fli1a:EGFP) y1, an EGFP-positive endothelial transgenic zebrafish line. Cancer cells were pretreated with CM and/or sLIFR as indicated. (H) Representative images of metastasis-like tumor structures (red) near vasculature (green) in zebrafish embryos. White arrows indicate dissemination sites. (I) Quantification of metastasis-like tumor structures on day 7 post-injection. Each dot represents one fish embryo. **P < 0.01, ***P < 0.001 (Bonferroni’s multiple comparison test). (J) Representative images of 3D spheroid assays. Macrophages (M□_GFP) were added to 1-day NPC-TW06_LifeAct-RFP spheroids (NPC_RFP) at a 1:1 effector-to-target ratio. Images were acquired 2 days post coculture. (K) Quantification of RFP-positive NPC spheroids using CellSens imaging software (Olympus). Each dot represents one spheroid. ****P < 0.000 (Bonferroni’s multiple comparison test).

### EBV product-containing exosomes induce immunosuppressive reprogramming of tumor and innate immune cells in the NPC microenvironment

To explore the impact of EBV-exosomes on the TME, we performed single-cell RNA sequencing (ScRNA-seq) on exosome-treated and control samples. Suspensions of cells freshly dissociated from NPC biopsy and treated with exosomes from HK1EBV cells or control medium for 2 hours were collected (Fig. 5A), yielding a total of 14,446 qualified cells. Subsequent cell typing and functional analyses identified epithelial cells, fibroblasts and seven major immune cell types comprising myeloid cells, T cells, natural killer (NK) cells, B cells, mast cells and neutrophiles (Fig. 5B). Cell type annotations were classified based on expression of canonical marker genes (Fig. 5C and Supplementary Table S4). Epithelial populations were further classified into non-cancer cells (83%) and cancer cells (17%) using inferCNV analysis (44). An increase in UMI (unique molecular identifier) counts of EBV-encoded LMP1 was detected in the exosome-treated group, indicating effective transfer of exosomal content (Fig. 5D). A gene set variation analysis (GSVA) revealed enhanced expression of genes associated with epithelial-mesenchymal transition (EMT), extracellular matrix (ECM) remodeling, IL6-JAK-STAT3 signaling, PD-1 signaling, and reactive oxygen species (ROS) pathways in cancer cells upon exosome treatment, whereas double-strand break (DSB) DNA repair, apoptosis, and chemokine signaling functions were suppressed (Fig. 5E). Consistent with this, PD-L1 and genes associated with EMT and ECM remodeling (*MMP14*, *MMP10*, *CTSV*, *VIM*, *AREG*, and *CTHRC1*) and the ROS pathway (*TXN*, *SOD1*, *PRDX1*, *PRDX6*, and *CAT*) were significantly upregulated in the exosome-treated group, whereas antigen-presentation genes (*TAPBP*, *CALR*, and *CANX*) were downregulated (Fig. 5F).

**Figure 5.**
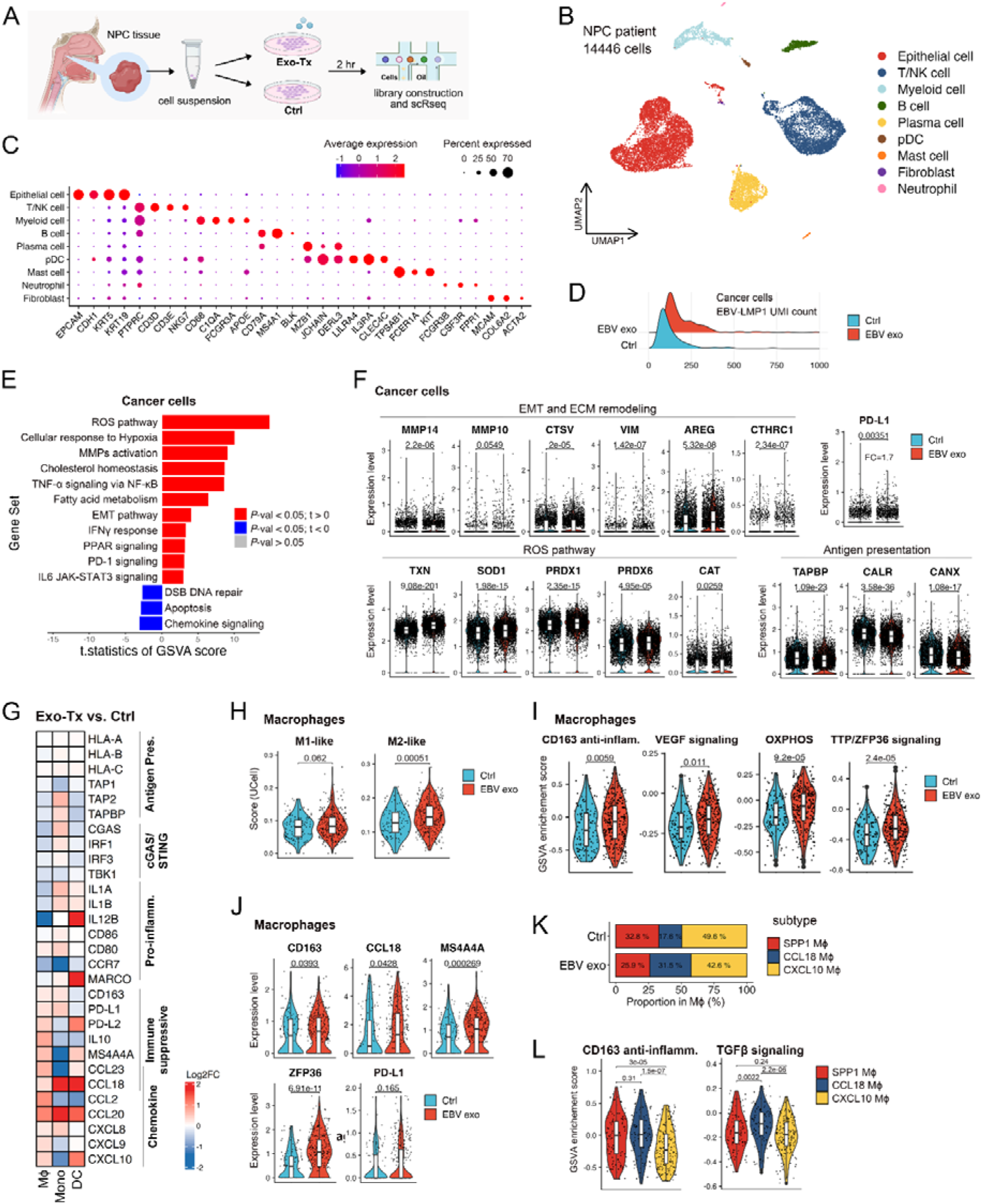
Exosome-mediated reprogramming of tumor and immune cells in the NPC TME. (A) Schematic of the ScRNA-seq experiment. (B) UMAP visualization of 14,446 qualified single cells grouped into nine major immune cell types. Each dot represents a single cell colored according to the annotated cell type. (C) Expression of representative marker genes used to define cell identities. Color scale indicates expression levels from blue (low) to red (high). (D) Ridge plot of UMI counts of EBV-encoded LMP1 in cancer cells. (E) GSVA enrichment scores comparing cancer cells treated with exosomes versus control samples. Red, enhanced pathways; blue, suppressed pathways. (F) Expression levels of PD-L1 and genes related to EMT, ECM remodeling, ROS, and antigen presentation in cancer cells (Mann-Whitney test). (G) Heatmap of representative DEGs involved in antigen presentation, cGAS-STING signaling, and pro-inflammatory, immunosuppressive, and chemokine pathways in macrophages, monocytes and dendritic cells. Values are presented as log_2_ fold changes (Log_2_FC) (exosome treatment versus control). Red, upregulation; blue, downregulation. (H) M1-like and M2-like signature scores of macrophages from control (blue) and exosome-treated (red) groups (Mann-Whitney U test). (I) GSVA enrichment scores for CD163-mediated anti-inflammatory, VEGF, OXPHOS, and TTP/ZFP36 signaling in macrophages (Mann-Whitney U test). (J) Expression of CD163, CCL18, MS4A4A, ZFP36, and PD-L1 in macrophages (Mann-Whitney U test). (K) Proportions of macrophage subtypes in control and exosome-treated groups. (L) GSVA enrichment scores for CD163-mediated anti-inflammatory and TGFB1 signaling in SPP1^+^, CCL18^+^, and CXCL10^+^ macrophage subsets.

In innate immune cells, including macrophages, monocytes and dendritic cells, exosome treatment led to altered expression of genes linked to antigen presentation, cGAS-STING signaling, immune suppression, inflammation, and chemokine activity (Fig. 5G). Macrophages showed a shift toward an M2-like phenotype, with increased M2 signature scores (Fig. 5H). Enrichment of CD163-mediated, anti-inflammatory VEGF, OXPHOS, and TTP/ZFP36 signaling further supported immune suppressive polarization (Fig. 5I). Upregulation of CD163, CCL18, MS4A4A, ZFP36, and PD-L1 expression in macrophages was observed following exosome treatment (Fig. 5J).

Subtype analysis revealed an exosome-mediated redistribution of macrophage populations, with increased proportions of CCL18^+^ macrophages (from 17.5% to 31.5%) and decreased SPP1^+^ (32.8% to 25.8%) and CXCL10^+^ (49.6% to 42.6%) macrophages (Fig. 5K). Among the three distinct TAM subpopulations CCL18-TAM exhibited relatively higher expression of FUCA1, CCL18, and ferroportin-associated genes SLC40A1 and SELENOP (SEPP1), which are known to promote tumorigenesis and M2-like polarization (45,46). SPP1-TAMs (SPP1, VCAN, INHBA) represented a pro-angiogenic population (47), whereas CXCL10-TAMs exerted pro-inflammatory effects by releasing CXCL10, CLEC10A, and antigen presentation-related CD1 molecules. The increased proportion of CCL18^+^ macrophages indicates an exosome-mediated immunosuppressive shift (45,46). In addition, among SPP1^+^, CCL18^+^ and CXCL10^+^ subsets, CD163-mediated anti-inflammatory and TGFB signaling were enhanced primarily in CCL18□ cells (Fig. 5L). These results demonstrate that tumor-derived exosomes modulate both tumor and immune cell phenotypes, promoting immunosuppression and tumor progression in the NPC microenvironment.

### EBV-exosomes enhance macrophage-T cell interactions and suppress T cell responses

Treatment with EBV product-containing exosomes broadly increased intercellular communication among major T cell subtypes and TAMs within the NPC TME. Cell–cell communication analysis further revealed exosome-mediated enhancement of ligand–receptor interactions between macrophages and T cells, including upregulated CCL20-CCR6 and CXCL9/10-CXCR3 axes (Fig. 6A).

**Figure 6.**
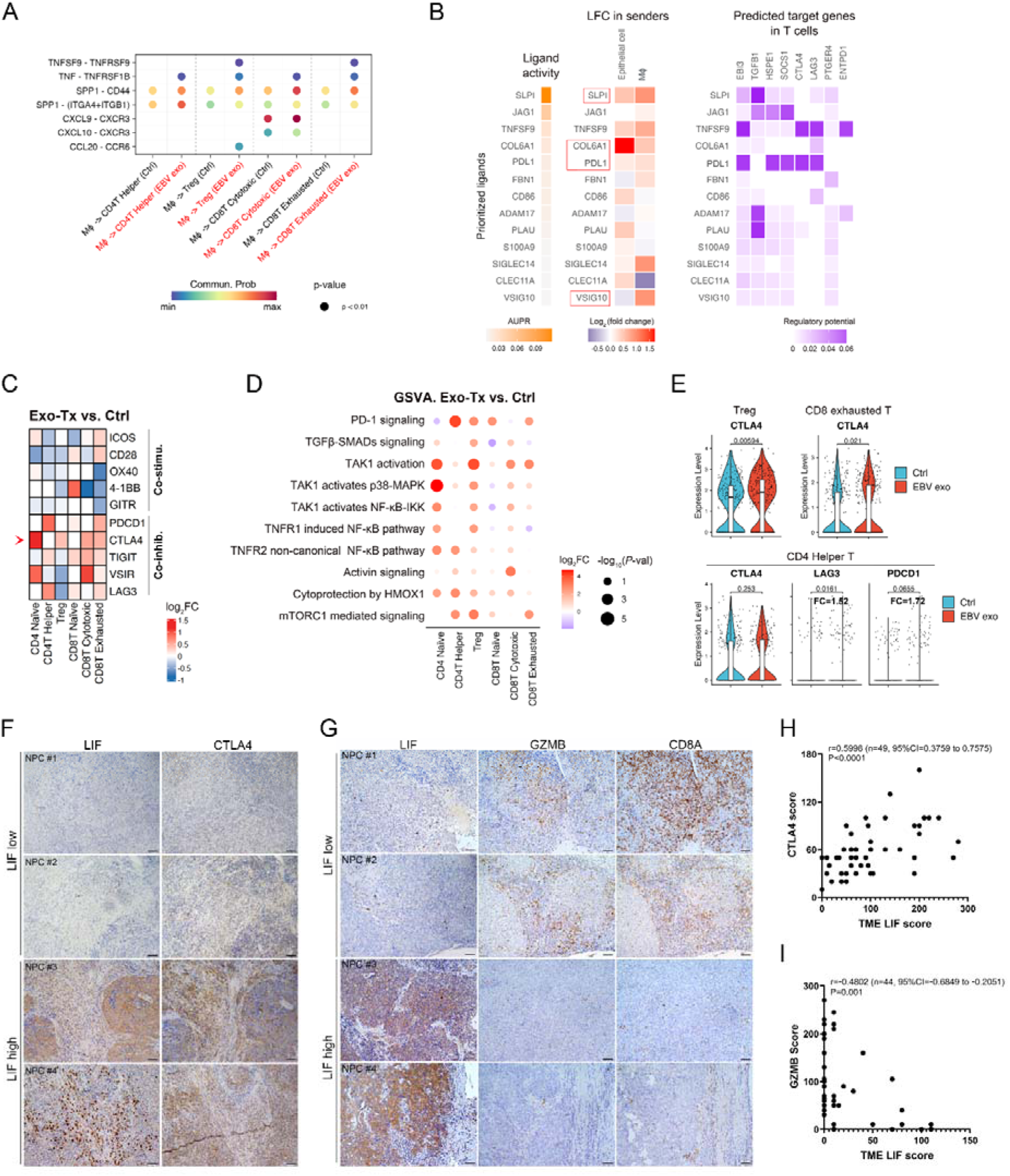
NPC-derived exosomes modulate T cell responses through TAM-mediated signaling. (A) Exosome-induced ligand-receptor interaction analysis by CellChat, with TAMs as sender cells and T cell subtypes as receivers. (B) NicheNet cell-cell communication analysis. Ligand and target gene activities were calculated based on the interactions between cancer cells and macrophages (ligand senders) and T cells (receiver). Left, AUPR scores of ligands. Middle, differential expression (Log_2_FC) of ligands in sender cells. Red, upregulated; blue, downregulated (Exo-Tx versus control). Right, predicted regulatory impact on target genes in T cells. Purple scale indicates relative regulatory potential. (C) Heatmap of representative DEGs related to T cell functions across major T cell subsets. Values are presented as Log_2_FC (Exo-Tx vs. control). Red, upregulated; blue, downregulated. (D) GSVA analysis of signaling pathways across major T cell subsets. Dot plots show t-statistics (color) and -log (P-value) (dot size) (paired t test). (E) Expression levels of CTLA4, LAG3, and PDCD1 in T cells from exosome-treated and control groups. (F) Representative IHC images of LIF and CTLA4 expression in consecutive NPC tumor sections. Scale bars, 50 μm. (G) Representative IHC images of LIF, GZMB, and CD8A in consecutive NPC biopsy section. Scale bars, 50 μm. (H and I) Correlation between LIF and CTLA4 expression scores (H), and LIF and GZMB/CD8A scores (I) in NPC TME. Spearman’s correlation coefficient (r), 95% confidence interval (CI), and *P*-values are indicated.

A NicheNet analysis identified macrophage/cancer-derived ligands that affect T cells. The top-ranked ligand, secretory leukocyte protease inhibitor (SLPI), is known to suppress inflammatory cytokines (12,13) (Fig. 6B). PD-L1 and COL6A1 were also implicated in T cell regulation. Exosome treatment induced T cell expression of co-inhibitory genes, including *CTLA4*, *PD-1*, *TIGIT* and *VSIR*, but suppressed co-stimulatory genes (*ICOS*, *CD28*, *OX40*, *4-1BB*, and *GITR*) (Fig. 6C). Moreover, results of GSVA revealed altered signaling pathways across T cell subsets following exosome exposure (Fig. 6D). Among these pathways were several canonical signaling pathways downstream of LIF and LMP1, including PD-1 signaling, TGFβ-SMAD signaling, NF-κB-associated cascades, and mTORC1 signaling (27,48,49). Among T cell subsets, Tregs and exhausted CD8□ T cells showed marked upregulation of CTLA4 (Fig. 6E).

IHC analyses showed enriched infiltration of CTLA4^+^ cells in NPC tumors with high LIF immunoreactivity in consecutive biopsy sections (Fig. 6F). Consistent with this, granzyme B (GZMB) and CD8A immunoreactivity were enhanced in the NPC TME with low LIF immunoreactivity and diminished in the TME with high LIF levels (Fig. 6G). Correlation analyses confirmed a positive correlation between LIF and CTLA4 expression (r = 0.5998, P < 0.0001; Fig. 6H) and an inverse correlation between LIF and GZMB/CD8A expression (r = -0.4802, *P* = 0.001; Fig. 6I) in consecutive NPC tissues. These findings further support the immunosuppressive function of LIF and the role of exosomal LMP1-LIF signaling in suppressing T cell responses via TAMs in the NPC TME.

Our findings collectively support a model in which EBV^+^ NPC-derived exosomes contribute to tumor progression by reprogramming macrophages toward an immunosuppressive phenotype. As schematically illustrated in Figure 7, tumor-derived exosomes containing LMP1 are taken up by macrophages, leading to NF-κB activation and upregulation of LIF. This LIF-driven signaling promotes polarization of TAMs and induces immunosuppressive crosstalk within the TME. The resulting macrophage-T cell interactions further dampen anti-tumor immunity and support cancer cell proliferation and dissemination. This model highlights the central role of the exosome-mediated LMP1-LIF axis in orchestrating immune suppression and facilitating tumor progression in NPC.

**Figure 7.**
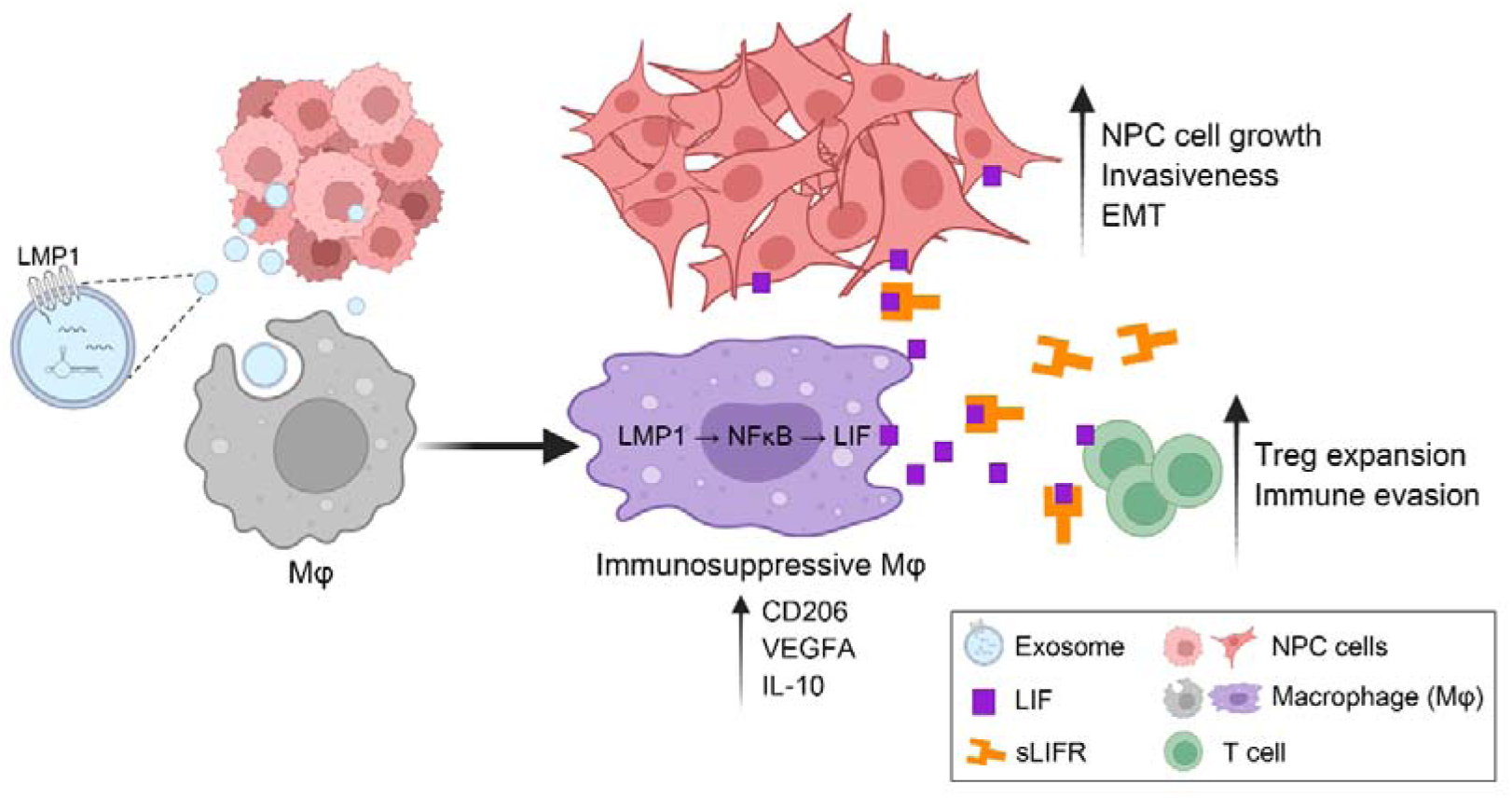
Schematic model for effects mediated by EBV product-containing exosomes in the NPC microenvironment.

## Discussion

In this study, we identified an exosomal LMP1-LIF axis that contributes to the immunosuppressive microenvironment of EBV-positive NPC. We showed that tumor-derived exosomes deliver LMP1 to macrophages, activating NF-κB signaling and inducing LIF expression. This process promoted macrophage polarization toward an immunosuppressive phenotype that interferes with T cell responses and facilitates tumor growth and dissemination. Our findings provide mechanistic insight into how EBV infection shapes the NPC immune landscape. While LIF has been associated with tumor progression in other malignancies, its regulation within macrophages in NPC was not well understood. By demonstrating that exosomal LMP1 drives LIF expression in TAMs, we identified a pathway that links viral oncogene activity to immune suppression. In our single-cell analyses, exosome exposure not only induced macrophage polarization but also coincided with increased CTLA4 and PD-1 signaling in T cells. This suggests that exosomal signaling operates in parallel with checkpoint pathways to reinforce an immunosuppressive state.

The broader implications extend beyond NPC. Exosome-mediated communication is a common feature of tumors, and EBV-associated malignancies such as Hodgkin lymphoma and EBV-positive gastric carcinoma may employ similar mechanisms. Understanding how viral products exploit exosome biology to reprogram immune cells could therefore provide a framework for studying immunosuppression in other virus-driven cancers.

These findings also raise questions relevant to therapeutic intervention. Although our data support LIF as a mediator of macrophage-induced tumor promotion, its inhibition alone may not be sufficient for therapeutic benefit. Instead, targeting the LMP1-LIF axis could be considered in combination with established immunotherapies such as checkpoint blockade. For example, suppressing exosome-induced LIF signaling might enhance T cell reinvigoration in the context of PD-1 or CTLA4 inhibition. Exploring such possibilities will require improved preclinical models.

Some limitations should also be acknowledged. The influence of other EBV-derived products and cytokines on this pathway remains to be defined, and the in vivo models we used cannot fully reproduce the complexity of human NPC. Establishing macrophage-exosome-NPC co-engraftment systems in mice remains technically difficult because of species-specific differences in exosome uptake and macrophage polarization. Moreover, our single-cell transcriptomic analyses reflect short-term exosome exposure rather than chronic interactions in the tumor microenvironment. Addressing these challenges will provide a more comprehensive understanding of how exosome-driven signaling sustains immunosuppression in NPC.

In conclusion, we showed that exosomal LMP1 promotes LIF-dependent macrophage polarization and contributes to immune suppression in NPC. By defining this previously unrecognized pathway of EBV-driven immune modulation, our study adds to the understanding of how tumor-derived exosomes shape the immune landscape and provides a foundation for future work exploring exosome biology in virus-associated cancers.

## Authors’ contributions

**Tzu-Tung Liu**: Performed single-cell experiments, bioinformatics analysis, IHC, immunofluorescence, western blotting, zebrafish experiment, and manuscript writing. **Yun-Hua Sui**: Conducted single-cell experiments, western blotting, and IHC. **Kai-Ping Chang**: Collected clinical samples and analyzed clinical data. **Fang-Yu Tsai:** Conducted single-cell transcriptomic analysis**. Ngan-Ming Tsang**: Performed IHC analysis. **Meng-Hsin Li** and **Hsuan-Chung Chen**: Conducted western blotting and IHC. **Chen-Han Huang:** Conducted TEM and provided technical support**. Ya-Wen Chen:** Provided resources. **Shih Sheng Jiang**: Conducted single-cell transcriptomic analysis, provided funding and resources. **Shu-Chen Liu**: Conceived the study, designed experiments, acquired funding, and revised the manuscript. All authors contributed to study conception, experimental design, data analysis, and interpretation.

## Acknowledgments

We appreciate the support from the Bioinformatics Core of the National Health Research Institutes, funded by the National Core Facility for Biopharmaceuticals (NCFB), National Science and Technology Council (NSTC113-2740-B-400-005).

## References

1. Wong KCW, Hui EP, Lo K-W, Lam WKJ, Johnson D, Li L, et al. Nasopharyngeal carcinoma: an evolving paradigm. Nature Reviews Clinical Oncology 2021;18:679–95

2. Jiang J, Ying H. Revealing the crosstalk between nasopharyngeal carcinoma and immune cells in the tumor microenvironment. Journal of Experimental & Clinical Cancer Research 2022;41:244

3. Shaofen H, Yunfan L, Rui D, Xiong L, Jie W, Lu W, et al. EBV-EBNA1 constructs an immunosuppressive microenvironment for nasopharyngeal carcinoma by promoting the chemoattraction of Treg cells. Journal for ImmunoTherapy of Cancer 2020;8:e001588

4. Li D, Qian L, Chen C, Shi M, Yu M, Hu M, et al. Down-regulation of MHC class II expression through inhibition of CIITA transcription by lytic transactivator Zta during Epstein-Barr virus reactivation. J Immunol 2009;182:1799–809

5. Yuan L, Zhong L, Krummenacher C, Zhao Q, Zhang X. Epstein–Barr virus-mediated immune evasion in tumor promotion. Trends in Immunology 2025;46:386–402

6. Cao Y, Xie L, Shi F, Tang M, Li Y, Hu J, et al. Targeting the signaling in Epstein-Barr virus-associated diseases: mechanism, regulation, and clinical study. Signal Transduction and Targeted Therapy 2021;6:15

7. Huang D, Song SJ, Wu ZZ, Wu W, Cui XY, Chen JN, et al. Epstein-Barr Virus-Induced VEGF and GM-CSF Drive Nasopharyngeal Carcinoma Metastasis via Recruitment and Activation of Macrophages. Cancer Res 2017;77:3591–604

8. Choi H, Lee H, Kim SR, Gho YS, Lee SK. Epstein-Barr virus-encoded microRNA BART15-3p promotes cell apoptosis partially by targeting BRUCE. J Virol 2013;87:8135–44

9. Lee PJ, Sui YH, Liu TT, Tsang NM, Huang CH, Lin TY, et al. Epstein-Barr viral product-containing exosomes promote fibrosis and nasopharyngeal carcinoma progression through activation of YAP1/FAPalpha signaling in fibroblasts. J Exp Clin Cancer Res 2022;41:254

10. Meckes DG, Jr., Shair KH, Marquitz AR, Kung CP, Edwards RH, Raab-Traub N. Human tumor virus utilizes exosomes for intercellular communication. Proc Natl Acad Sci U S A 2010;107:20370–5

11. Metcalf D. The unsolved enigmas of leukemia inhibitory factor. Stem Cells 2003;21:5–14

12. Auernhammer CJ, Melmed S. Leukemia-inhibitory factor-neuroimmune modulator of endocrine function. Endocr Rev 2000;21:313–45

13. Burdon T, Smith A, Savatier P. Signalling, cell cycle and pluripotency in embryonic stem cells. Trends Cell Biol 2002;12:432–8

14. Pera MF, Tam PP. Extrinsic regulation of pluripotent stem cells. Nature 2010;465:713–20

15. Zouein FA, Kurdi M, Booz GW. LIF and the heart: just another brick in the wall? Eur Cytokine Netw 2013;24:11–9

16. Mathieu ME, Saucourt C, Mournetas V, Gauthereau X, Theze N, Praloran V, et al. LIF-dependent signaling: new pieces in the Lego. Stem Cell Rev 2012;8:1–15

17. Trouillas M, Saucourt C, Guillotin B, Gauthereau X, Taupin JL, Moreau JF, et al. The LIF cytokine: towards adulthood. Eur Cytokine Netw 2009;20:51–62

18. Liu SC, Tsang NM, Chiang WC, Chang KP, Hsueh C, Liang Y, et al. Leukemia inhibitory factor promotes nasopharyngeal carcinoma progression and radioresistance. J Clin Invest 2013;123:5269–83

19. Liu SC, Tsang NM, Chiang WC, Chang KP, Hsueh C, Liang Y, et al. Leukemia inhibitory factor promotes nasopharyngeal carcinoma progression and radioresistance. J Clin Invest 2013;123:5269–83

20. Metcalfe SM. LIF in the regulation of T-cell fate and as a potential therapeutic. Genes Immun 2011;12:157–68

21. Cao W, Yang Y, Wang Z, Liu A, Fang L, Wu F, et al. Leukemia inhibitory factor inhibits T helper 17 cell differentiation and confers treatment effects of neural progenitor cell therapy in autoimmune disease. Immunity 2011;35:273–84

22. Gao W, Thompson L, Zhou Q, Putheti P, Fahmy TM, Strom TB, et al. Treg versus Th17 lymphocyte lineages are cross-regulated by LIF versus IL-6. Cell Cycle 2009;8:1444–50

23. Duluc D, Delneste Y, Tan F, Moles MP, Grimaud L, Lenoir J, et al. Tumor-associated leukemia inhibitory factor and IL-6 skew monocyte differentiation into tumor-associated macrophage-like cells. Blood 2007;110:4319–30

24. Jeannin P, Duluc D, Delneste Y. IL-6 and leukemia-inhibitory factor are involved in the generation of tumor-associated macrophage: regulation by IFN-gamma. Immunotherapy 2011;3:23–6

25. Won H, Moreira D, Gao C, Duttagupta P, Zhao X, Manuel E, et al. TLR9 expression and secretion of LIF by prostate cancer cells stimulates accumulation and activity of polymorphonuclear MDSCs. J Leukoc Biol 2017;102:423–36

26. Pascual-Garcia M, Bonfill-Teixidor E, Planas-Rigol E, Rubio-Perez C, Iurlaro R, Arias A, et al. LIF regulates CXCL9 in tumor-associated macrophages and prevents CD8(+) T cell tumor-infiltration impairing anti-PD1 therapy. Nat Commun 2019;10:2416

27. Liu S-C, Tsang N-M, Chiang W-C, Chang K-P, Hsueh C, Liang Y, et al. Leukemia inhibitory factor promotes nasopharyngeal carcinoma progression and radioresistance. The Journal of Clinical Investigation 2013;123:5269–83

28. Lo AK, Lo KW, Tsao SW, Wong HL, Hui JW, To KF, et al. Epstein-Barr virus infection alters cellular signal cascades in human nasopharyngeal epithelial cells. Neoplasia 2006;8:173–80

29. Miller G, Shope T, Lisco H, Stitt D, Lipman M. Epstein-Barr virus: transformation, cytopathic changes, and viral antigens in squirrel monkey and marmoset leukocytes. Proc Natl Acad Sci U S A 1972;69:383–7

30. Liao SK, Perng YP, Shen YC, Chung PJ, Chang YS, Wang CH. Chromosomal abnormalities of a new nasopharyngeal carcinoma cell line (NPC-BM1) derived from a bone marrow metastatic lesion. Cancer Genet Cytogenet 1998;103:52–8

31. Lin CT, Chan WY, Chen W, Huang HM, Wu HC, Hsu MM, et al. Characterization of seven newly established nasopharyngeal carcinoma cell lines. Lab Invest 1993;68:716–27

32. Kong AT, Leprevost FV, Avtonomov DM, Mellacheruvu D, Nesvizhskii AI. MSFragger: ultrafast and comprehensive peptide identification in mass spectrometry-based proteomics. Nature Methods 2017;14:513–20

33. Liu SC, Hsu T, Chang YS, Chung AK, Jiang SS, OuYang CN, et al. Cytoplasmic LIF reprograms invasive mode to enhance NPC dissemination through modulating YAP1-FAK/PXN signaling. Nat Commun 2018;9:5105

34. Chen CC, Chen LC, Liang Y, Tsang NM, Chang YS. Epstein-Barr virus latent membrane protein 1 induces the chemotherapeutic target, thymidine phosphorylase, via NF-kappaB and p38 MAPK pathways. Cell Signal 2010;22:1132–42

35. Liu HP, Wu CC, Chang YS. PRA1 promotes the intracellular trafficking and NF-kappaB signaling of EBV latent membrane protein 1. Embo j 2006;25:4120–30

36. Hao Y, Stuart T, Kowalski MH, Choudhary S, Hoffman P, Hartman A, et al. Dictionary learning for integrative, multimodal and scalable single-cell analysis. Nat Biotechnol 2024;42:293–304

37. McGinnis CS, Murrow LM, Gartner ZJ. DoubletFinder: Doublet Detection in Single-Cell RNA Sequencing Data Using Artificial Nearest Neighbors. Cell Systems 2019;8:329–37.e4

38. Coulton A, Murai J, Qian D, Thakkar K, Lewis CE, Litchfield K. Using a pan-cancer atlas to investigate tumour associated macrophages as regulators of immunotherapy response. Nature Communications 2024;15:5665

39. Hänzelmann S, Castelo R, Guinney J. GSVA: gene set variation analysis for microarray and RNA-Seq data. BMC Bioinformatics 2013;14:7

40. Browaeys R, Saelens W, Saeys Y. NicheNet: modeling intercellular communication by linking ligands to target genes. Nat Methods 2020;17:159–62

41. Viswanadhapalli S, Dileep KV, Zhang KYJ, Nair HB, Vadlamudi RK. Targeting LIF/LIFR signaling in cancer. Genes Dis 2022;9:973–80

42. Wang J, Chang CY, Yang X, Zhou F, Liu J, Feng Z, et al. Leukemia inhibitory factor, a double-edged sword with therapeutic implications in human diseases. Mol Ther 2023;31:331–43

43. Takada K, Horinouchi K, Ono Y, Aya T, Osato T, Takahashi M, et al. An Epstein-Barr virus-producer line Akata: establishment of the cell line and analysis of viral DNA. Virus Genes 1991;5:147–56

44. Patel AP, Tirosh I, Trombetta JJ, Shalek AK, Gillespie SM, Wakimoto H, et al. Single-cell RNA-seq highlights intratumoral heterogeneity in primary glioblastoma. Science 2014;344:1396–401

45. Barrett CW, Reddy VK, Short SP, Motley AK, Lintel MK, Bradley AM, et al. Selenoprotein P influences colitis-induced tumorigenesis by mediating stemness and oxidative damage. J Clin Invest 2015;125:2646–60

46. Qian J, Olbrecht S, Boeckx B, Vos H, Laoui D, Etlioglu E, et al. A pan-cancer blueprint of the heterogeneous tumor microenvironment revealed by single-cell profiling. Cell Research 2020;30:745–62

47. Cheng S, Li Z, Gao R, Xing B, Gao Y, Yang Y, et al. A pan-cancer single-cell transcriptional atlas of tumor infiltrating myeloid cells. Cell 2021;184:792–809.e23

48. Luo Y, Liu Y, Wang C, Gan R. Signaling pathways of EBV-induced oncogenesis. Cancer Cell International 2021;21:93

49. Lo AKF, Dawson CW, Lo KW, Yu Y, Young LS. Upregulation of Id1 by Epstein-Barr Virus-encoded LMP1 confers resistance to TGFβ-mediated growth inhibition. Molecular Cancer 2010;9:155

